# Modeling Masticatory Myalgia to Headache-like Referred Pain Triggers Local Gene Plasticity at Referred Pain Sites

**DOI:** 10.1101/2025.03.26.645551

**Authors:** Anahit H. Hovhannisyan, Jessica Aldape, Jennifer M. Mecklenburg, Jessie Alfaro, Yi Zou, Zhao Lai, Wei Guo, Jiale Yang, Malin Ernberg, M. Danilo Boada, Ke Ren, Armen N. Akopian

**Author notes:** **Corresponding authors:** Armen N. Akopian, University of Texas Health Science Center @ San Antonio, 7703 Floyd Curl Drive, San Antonio, TX 78229-3900.

## Abstract

Patients with myofascial pain in the head and neck area report widespread and referred pain, including headache. Existing preclinical models fail to replicate this clinical phenotype; therefore, we aimed to develop animal models mimicking referred pain phenomenon and investigate whether referred pain leads to gene plasticity at the referred sites. We modeled masticatory myalgia by stimulation of either the masseter (MM) or temporal muscle (TM) in mice. MM and TM were stimulated with a single high-dose injection of Collagenase-type II (Col), repetitive low-dose Col injections, repetitive gentle MM stimulation, or single or repetitive forceful mouth opening. Referred pain was assessed by measuring mechanical hypersensitivity in the periorbital area (representing headache-like behavior) and another masticatory muscle. Stimulation of the MM, whether through single or repetitive Col injections or mouth opening, produced inconsistent, short-lasting (1-2 days) headache-like behavior in both males and females. In contrast, stimulation of the TM, using different paradigms, triggered mechanical hypersensitivity in both the MM and the periorbital area. Referred headache-like behavior lasted longer in females compared to males, while referred myalgia in the MM was pronouncer in males. The referred pain in the MM and periorbital areas triggered by TM stimulation was associated with significant gene plasticity in the MM and dura mater. Transcriptional changes in the MM following Col injection into the TM resembled those observed after direct MM injections. Presented data imply that referred pain modeled by TM stimulation could be accounted by nociceptive signaling from multiple local sites involved in this referred pain network.

## Introduction

Chronic musculoskeletal pain conditions, such as myogenous temporomandibular disorders (TMD-M), have multifactorial backgrounds that have a significant impact on the individual and society [1; 36]. TMD-M is classified to local myalgia (MYA) or myofascial pain with pain referral (MFP) according to the diagnostic criteria for temporomandibular disorders (DC/TMD) [5; 39]. MYA is characterized by local pain, which is frequently patients’ main complaint. MFP patients report widespread and referred pain [5; 29]. Thus, MFP patients often report symptoms such as headache, and pain within the temporomandibular joint (TMJ) and neck [31].

The mechanisms for referred pain are unclear, but several possibilities have been suggested. Referred pain could be due to sensitization of second-order brain stem neurons, which are convergence points for sensory neuron afferents innervating different parts of the head and neck area, which will lead to expansion of receptive fields [10; 19]. Referred pain could also be a sign of central sensitization due to inhibition of a descending pain pathway [13; 16; 42]. Local mechanisms could also contribute to referred pain, since stronger, longer, and more painful stimulations of muscles produced more referred pain [13; 42]. Moreover, local anesthetics reduced referred pain [42]. Overall, understanding intricate and complex mechanisms of referred pain could highly benefit from studies on preclinical models for referred pain. Such animal studies will produce essential information and data on gene and cell plasticity in the peripheral and central portions of the nociceptive pathways.

In this respect, the overarching goal of this study was to model head and neck area referral pain in mice. Using such modeling, we investigated whether masticatory myalgia referred headache-like behavior relies on which muscle (i.e masseter versus temporalis) elicits pain; whether referred pain was sex-dependent and whether headache-like behavior and wide-spread masticatory myalgia elicited from a temporalis muscle (TM) led to local gene plasticity at the referred pain sites, such as dura mater and masseter muscle (MM).

## Methods

### Ethical Approval

This study adheres to the ARRIVE 2.0 guidelines [38]. All animal care and experimental procedures complied with the United States Public Health Service Policy on Humane Care and Use of Laboratory Animals, the Guide for the Care and Use of Laboratory Animals. We followed guidelines from the National Institutes of Health (NIH) and the Society for Neuroscience (SfN) to minimize the number of animals used and their suffering. All mice were euthanized by cervical dislocation under isoflurane anesthesia followed immediately by decapitation. Isoflurane ensured that the mice were unconscious, while dislocation provided for expedient euthanasia. Decapitation assured death. This procedure minimized animal distress, and recommended by the AVMA Guidelines for the Euthanasia of Animals. All animal experiments conformed to protocols approved by the University Texas Health Science Center at San Antonio (UTHSCSA) and the University of Maryland Institutional Animal Care and Use Committee (IACUC). The IACUC protocol numbers in UTHSCSA are 20190114AR, 20220064AR and 20220069AR.

### Key reagents and mouse lines

Mice were housed under controlled conditions (≈22°C), relative humidity 40-60%, and a 12-h light-dark cycle with lights on at 7:00 AM. Food and water were available *ad libitum* in their home cages. Experiments were conducted on wild-type adult (2-4-months-old) C57Bl/6 male and female mice purchased from Jackson Laboratory (Bar Harbor, ME). Crude collagenase type II (Col; >125 units per mg; Cat: LS004214) preparations were acquired from Worthington (Lakewood, NJ). Colibri Retractors (item No: 17000-03) were acquired from Fine Science Tools (Foster City, CA). A vibrator was Point Relief massager (Fabrication Enterprises, Elmsford, NY).

### Masticatory myalgia models

In this study, we have used the following masticatory myalgia models:

#### Col MM model [26]

Left side MM was injected with 10μl Col. Injection was performed once or repetitively every 4 days. Col dosage was 0.2U, 0.5U or 10U. These injections were performed under isoflurane anesthesia. The area of injection was swabbed with 70% alcohol beforehand. The site for injection was identified by palpating the zygomatic arch and the body of the mandible. A 28-gauge needle was inserted into the point inferior to the posterior third of the zygomatic arch and midway between the zygomatic arch and the body of the mandible. The needle was advanced in an anterior direction in about 3 mm. After a gentle aspiration, solutions were injected. PBS served as vehicle control.

#### Col MM with repetitive gentle MM stimulation model (adapted from [26])

Left side MM was injected with 10μl Col. Dosages of Col were 0.2U or 1U. Gentle repetitive MM stimulations (2×90 sec) were applied every day by Point Relief vibrator using a sharp rubber insert. Gentle repetitive MM stimulations were always applied after measurements of mechanical hypersensitivity. Col injections and vibration stimuli were performed under isoflurane anesthesia.

#### Col TM model (adapted from [26])

Left side TM was injected with 10μl Col. Injection was performed once or repetitively every 4 days. Col dosage was 0.2U, 0.5U or 10U. These injections were performed under isoflurane anesthesia. Injection was performed into a middle part of TM. The syringe’s needle was directed slightly parallel to the head. A needle penetrated the muscle and the injected solution did not generate “bubbles” or air pockets under the skin.

#### Mouth stretch model (adapted from [34])

Mice mouths were forcibly stretched by placement of a 3 cm Colibri Retractor along top and bottom incisors following anesthetization. Mouth stretching lasted 5-, 15-, or 45-min. Stretching was performed once or repeated every 4^th^ day. Mice were anesthetized by injecting 40 μl of a cocktail of 156μl ketamine + 250μl dexmedetomidine into one thigh muscle. Once animals were anesthetized, forceps were used to open mouth and put the refractor inside the mouth where the teeth are holding it in place. Mice were woken up by an intramuscular injection of 40µl of Revertidine.

### Measurements of orofacial pain and headache-like behavior

#### Measurement of mechanical thresholds with von Frey filaments

At UTHSCSA, the mechanical hypersensitivity of the orofacial region was assessed as previously described [23; 30]. Briefly, mechanical hypersensitivity was assessed in several locations of an orofacial region, including the left side of the MM [20; 26], TMJ [26] and periorbital areas [3] (see schematic on *Fig 1A*). To avoid any anticipatory movement from mice during von Frey application, the mice were habituated to small cages for 3 days, followed by habituation to a designated area on a table and experimenter’s hand for an additional 3 days. During the last 3 days, mice were also habituated to von Frey filament (0.07-0.6g) stimulations in different orofacial areas. Measurements with von Frey were conducted on the animals kept in hands or freely moving on table. Habituated naive mice that did not respond to 0.6g von Frey filaments, which is considered baseline, were selected for the experimental procedure [2; 3; 26]. Experimental mice were probed with 0.008-0.6g filaments to the orofacial areas using an up-down approach [3; 20; 26]. Intervals between von Frey filament applications were varied but >30 sec. Mechanical thresholds were assessed and calculated as previously described [2; 20; 26].

**Figure 1.**
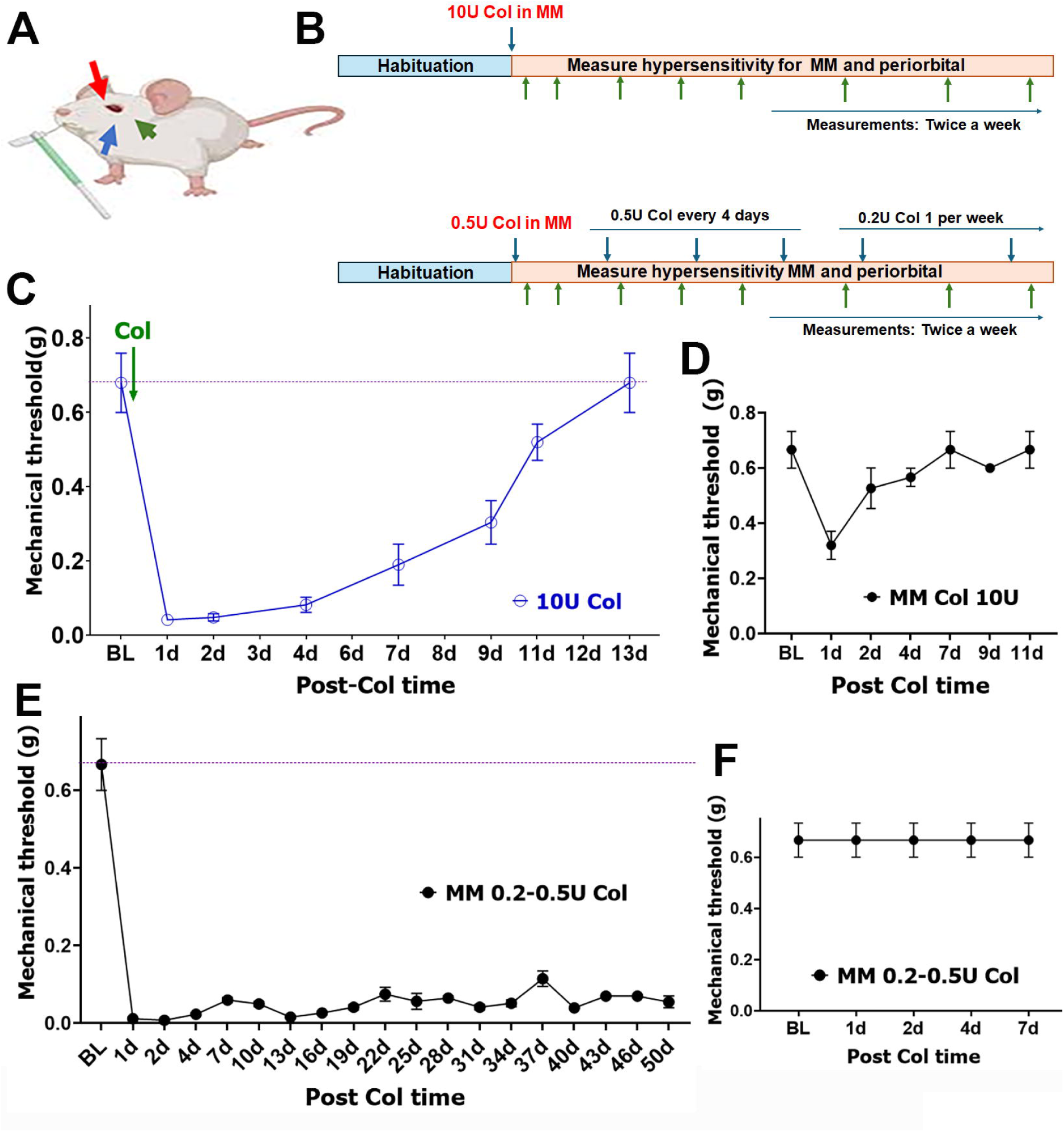
Referred pain after modeling masticatory myalgia by stimulating masseter muscle with collagenase type II. **(A)** Locations of head and neck areas in mice for von Frey applications and measurements of mechanical hypersensitivity. The blue arrow shows a mandible area. The green arrow indicate an area over masseter muscle (MM). The red arrow shows a periorbital area. **(B)** Schematic of paradigms for MM stimulations with single 10U collagenase type II (Col; upper panel) injection and repetitive 0.2-0.5U Col injections (bottom panel). **(C)** The development of masticatory myalgia at an area over MM after single 10U Col injection in male mice. **(D)** The development of periorbital mechanical hypersensitivity after single 10U Col injection into MM of male mice. **(E)** The development of masticatory myalgia at an area over MM after repetitive 0.2-0.5U Col injections in male mice. **(F)** The development of periorbital mechanical hypersensitivity after repetitive 0.2-0.5U Col injections into MM of male mice.

#### Measurement of mechanical sensitivity

At the University of Maryland, the mechanical sensitivity test of the orofacial region was performed as previously described [20]. Briefly, mice were habituated to be held entirely in the experimenter’s hand wearing a leather work glove. A series of calibrated von Frey filaments were applied to the skin above the injured MM. An active withdrawal of the head from the probing filament was defined as a response. Each von Frey filament was applied 5 times at intervals of 5-10 seconds. The response frequencies [(the number of responses/number of stimuli) X100%] to a range of von Frey filament forces were determined and a stimulus-response frequency (S-R) curve was plotted. After non-linear regression analysis, an EF_50_ value, defined as the effective von Frey filament force (g) that produces a 50% response frequency, was derived from the S-R curve (Prism, GraphPad) [19]. The injection of Col into the MM resulted in a leftward shift of the S-R curve and a reduction of EF_50_, suggesting the development of mechanical hypersensitivity.

#### Measurement of conditioned place avoidance behavior (CPA)

At the University of Maryland, for the CPA test [17; 25], animals were placed within a 42 (L) x 18 (W) x 18 (H) cm Plexiglass chamber. One half of the chamber was made with white plates (light area) and the other half was made with black plates (dark area). During the test, animals were allowed unrestricted movement throughout the test chambers for the 30-min test period. Mice normally prefer to stay in the dark area. Testing began immediately with suprathreshold mechanical stimulation (117 mN von Frey monofilament) applied to the facial skin at 15-sec intervals. The mechanical stimulus was applied to the skin above the injured muscle when the animal was within the preferred dark area and to the facial skin on the non-injured side when the animal was within the non-preferred light side. Based on the location of the animal at each 15-sec interval, the mean percentage of time spent in the black (dark) or white (light) chamber was calculated. Staying in the preferred dark area was associated with an aversive and painful stimulus after Col injection and the injured mouse tended to leave the dark side to avoid the aversive stimulus. An increase in the time spent in the light area suggests an increase in stimulus aversiveness.

### RNA seq transcriptomic data generation, analyses, and statistics

MM was dissected from left side. Dissected MM included deep, middle and superficial parts, and maximally excluded tendons attaching MM to zygomatic bone and mandible. Frozen and sliced MM as previously described [27] and freshly dissected entire dura matter were homogenized in Rn-easy solution (Qiagen) using a Bead Mill Homogenizer (Omni International, Kennesaw, GA). Extraction of RNA, RNA quality and integrity control, cDNA library preparation with oligo-dT primers followed the Illumina TruSeq stranded mRNA sample preparation guide (Illumina, San Diego, CA), and sequencing procedure with Illumina HiSeq 3000 platform with 30-50×10^6^ bp reading depth were previously described in detail [20; 26; 33]. Total RNA quality of between 7 and 9 were used in RNA-seq experiments. This is a critical point, since power analysis is based on these numbers for quality of total RNA.

Post-sequencing de-multiplexing with CASAVA, the combined raw reads were aligned to mouse genome build UCSC/mm10 using TopHat2 default settings and differentially expressed genes (DEGs) were identified using DESeq2 after performing median normalization, also previously described in detail [20; 26; 33].

Quality control statistical analysis of outliers, intergroup variability, distribution levels, PCA, and hierarchical clustering analysis were performed to statistically validate the experimental data. Multiple correction test was performed with the Benjamini-Hochberg procedure and adjusted p-value (Padj) was generated. If not specified in the text, criteria for selecting DEGs were expression levels with RPKM>1, fold-change (FC)>1.5, and statistically significant DEGs with Padj<0.05. Venn diagrams were generated using https://bioinfogp.cnb.csic.es/tools/venny/. Genes were clustered according to biological processes using the PANTHER software (http://www.pantherdb.org/).

### Statistical Analyses

Sample size calculations were based on our previous experience in RNA-seq and behavioral experiments on linear and inbred C57-black mice [2; 3; 20; 26; 32; 33]. Accordingly, in behavioral experiments, we estimate that 6-8 mice will achieve > 80% statistical power to achieve p<0.05 difference in measurement of hypersensitivity. For bulk RNA-seq, a sample size of 3-4 mice per group, using false discovery rate (FDR) of 0.05, will achieve > 80% statistical power to detect a two-fold change in gene expression with an estimated standard deviation of 0.5. We also assume 1% of genes are differentially expressed in the system, or 100 differentially expressed genes among the typical 10,000 expressed genes in tissues profiled. Using Scotty [9], we estimate that 40 million reads/samples are needed.

GraphPad Prism 8.0 (GraphPad, La Jolla, CA) was used for statistical analysis. Data are presented as mean ± standard error of mean (SEM). Differences between groups were assessed by chi-square analysis with Fisher’s exact test, unpaired t-test, and 1-way or 2-way ANOVA with Bonferroni’s post-hoc test. A difference is accepted as statistically significant when p<0.05. Interaction F ratios, and the associated p values are reported.

## Results

The objectives of this study were to evaluate whether modeled masticatory myalgia via different paradigms generated referred pain in adjusted masticatory muscles and headache-like behavior; and to examine whether referred pain is accompanied with gene plasticity in local tissues to which referred pain was extended. Distinct paradigms for modeling masticatory myalgia involved stimulation of different facial muscles (MM and TM), elicit muscle stimulation by mouth stretch, vibration and/or collagenase type-II (Col) intramuscular injections.

### Referred pain after eliciting masticatory myalgia from masseter muscle

Mechanical hypersensitivity representing masticatory myalgia was evaluated by probing orofacial area over MM (close to TMJ area) with von Frey (*Fig 1A*; green arrow), while headache-like behavior was assessed by probing von Frey at periorbital area (*Fig 1A*; red arrow) [3; 20; 26]. Experimental schematic showed on *Fig 1B* (upper panel). After habituation, a single 10U Col injection was administered, and myalgia and headache-like behavior was monitored longitudinally in male mice. Myalgia lasted up-to 2 weeks as it was reported before (*Fig 1C*) [26]. Headache-like behavior for this experimental paradigm was registered only in 30% of tested animals, was mild and lasted up to 4 days (*Fig 1D*). These data indicated that stimulation of MM by a single high-dose injection of Col generated pronounced myalgia, but referred headache-like pain was mild and not revealed in all animals.

Masticatory myalgia usually does not show high inflammatory responses in facial muscles [35; 47]. Unlike high dosage Col (10U), low-dose stimulation (0.1-0.2U) did not produce inflammatory response in MM [26]. Hence, we have modeled myalgia by repetitive application of 0.2-0.5U Col (*Fig 1B*; lower panel). This model produced persistent myalgia as long as Col was repetitively applied (*Fig 1E*), while headache-like behavior was not detected for entire length of experiments (*Fig 1F* shows first 7 days as an example). These data showed that stimulation of MM by low-dose injection of Col generated persistent myalgia, but referred pain was absent in all evaluated male mice.

The literature reports inconsistent epidemiological values concerning TMD-M prevalence. However, there is an agreement that women are 2-4 times more likely to experience masticatory activity-induced pain than men [8; 22; 41]. Accordingly, we conducted an independent study (at the University of Maryland) to examine whether intramuscular Col treatment produces sex-dependent masticatory myalgia. Col induced mechanical hypersensitivity in a dose-dependent fashion in both males (2-way ANOVA; interaction F (21, 192) = 6.540, P<0.0001; n=7, *Fig 2A*) and females (2-way ANOVA; interaction F (21, 192) = 8.013, P<0.0001, n=7, *Fig 2B*). At a lower dose of 0.2U Col, mechanical hypersensitivity was statistically similar in females versus males (2-way ANOVA; interaction F (7, 96) = 0.4346; P=0.8782; n=7; *Fig 2C*). An increase of Col dose to 10U generated sex differences in mechanical hypersensitivity with myalgia lasting slightly longer in males compared to females (2-way ANOVA; interaction F (7, 96) = 4.615; P=0.0002; n=7; *Fig 2D*). Patients with TMD-M often report spontaneous pain [18; 37]. Here, we employed the conditioned place avoidance (CPA) assay to evaluate this component in female and male mice [17]. Male mice injected with 10U of Col into MM exhibited robust CPA behavior at 10d that remained significant at day 21 (1-way ANOVA compared to baseline (BL); F (2, 18) = 113.9; P<0.0001; n=7; *Fig 2E*). Baseline responses in females were like males (un-published data; [17]). Administration of 10U Col into the MM of females produced CPA behavior at 10d, which was like behavior observed in males (*Fig 2F vs. 2E*). By 21d post-Col, spontaneous pain was slightly reduced in females versus males (28.75±4.09 in males versus 22.14±2.91 in females; *Fig 2F vs. 2E*). Overall, Col-induced masticatory myalgia after stimulation of MM was slightly more pronounced in males than females.

**Figure 2.**
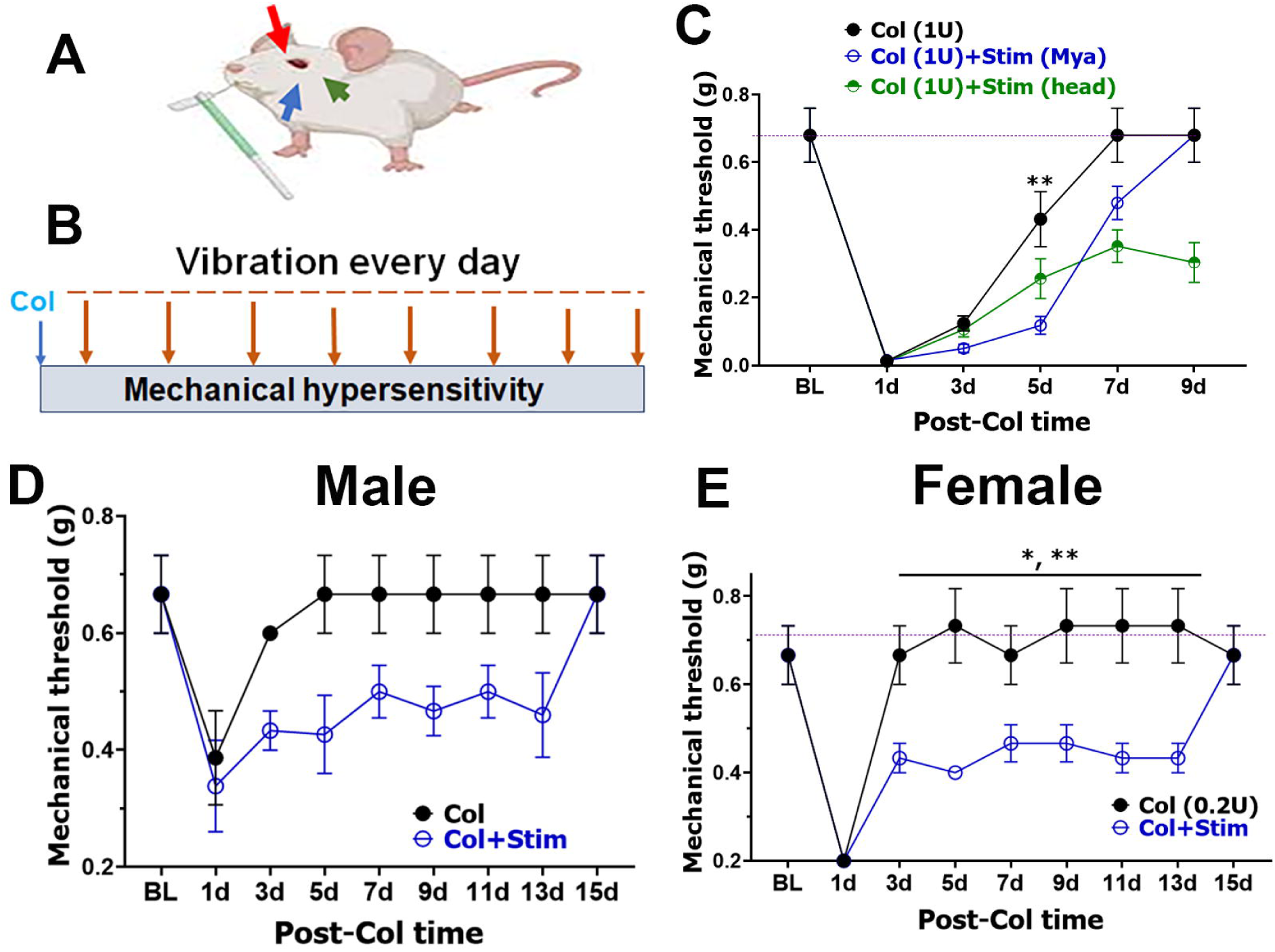
Masticatory myalgia elicited by Col treatment of MM in females and males. (**A, B**) Col-induced mechanical hypersensitivity in male (*panel A*) and female (*panel B*) mice was measured by von Frey filaments. Col injection doses are indicated. Statistical analysis is a 2-way ANOVA with Bonferroni’s post-hoc test calculated relative to Veh (PBS) (* p<0.05; ** p<0.01; *** p<0.001; ^#^ p<0.0001; n=7). (**C, D**) Col-induced mechanical hypersensitivity in male versus female mice for 0.2U Col (*panel C)* and 10U Col (*panel D*). Statistical analysis is a 2-way ANOVA with Bonferroni’s post-hoc test (** p<0.01; ^#^ p<0.0001; n=7). (**E, F**) Col (10U)-induced hypersensitivity in male (*panel E*) and female (*panel F*) mice was measured by conditioned place avoidance (CPA) behavior. Statistical analysis is a 1-way ANOVA with Bonferroni’s post-hoc test calculated relatively to pre-Col reading (NS – non-significant; * p<0.05; ^#^ p<0.0001; n=7).

It is presumed that masticatory myalgia develops due to muscle ischemia following consistent and strong or mild repetitive physical overloading of facial muscles [15; 28]. We have mimicked mild repetitive overload of muscle by applying daily vibration over MM at points indicated by blue and green arrows (*Fig 3A*). Such repetitive mild vibration alone did not induce either myalgia or headache. However, this type of daily mild stimulation of MM coupled with lower dosage Col (1 U) injection prolonged masticatory myalgia and generated headache-like behavior (F (10, 72) = 4.059; P=0.0002; n=5; *Fig 3B, 3C*). Headache-like behavior lasted as long as vibration-stimulation continued. This effect was similar in male and female mice (*Figs 3D, 3E*). Nevertheless, effect of daily vibration over MM on generation of headache-like pain reached statistical significance only for females, when Col dosage was reduced to 0.2U (for males F (8, 90) = 1.095; P=0.37; n=6; and for females F (8, 89) = 2.707; P=0.01; n=6; *Figs 3E, 3F*). This set of data implied that to induce referred pain (headache) by eliciting stimulation of MM, combination of stimuli such as single low-dose Col injection and repetitive vibration over MM are required. Moreover, certain level of sex dimorphism was noted for the described myalgia paradigm leading to referred pain.

**Figure 3.**
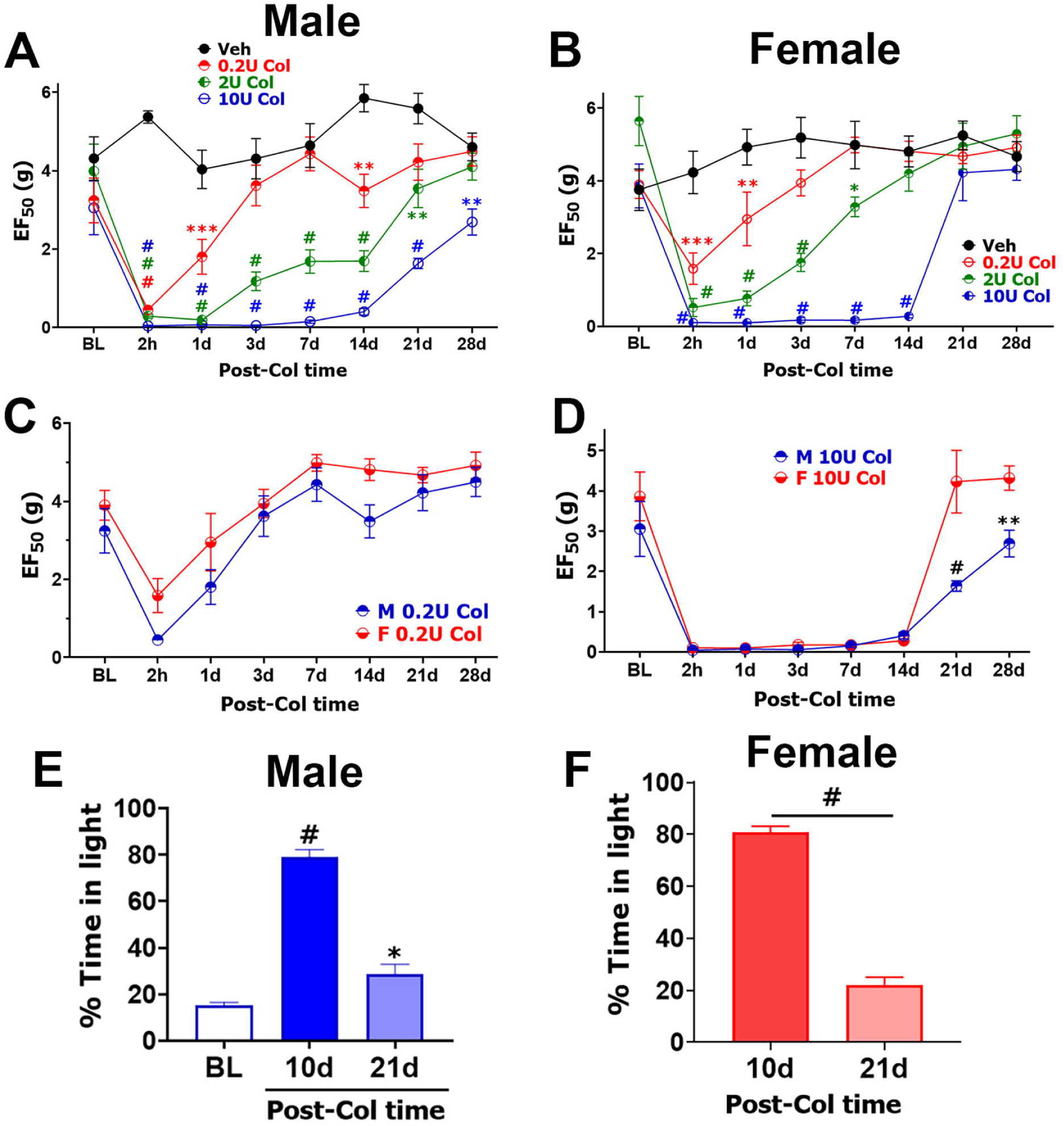
Referred pain after modeling masticatory myalgia by stimulating MM with Col and repetitive vibrations over MM. **(A)** Locations of head and neck areas in mice for von Frey applications and measurements of mechanical hypersensitivity. The blue arrow shows a mandible area. The green arrow indicate an area over MM. The red arrow shows a periorbital area. **(B)** Schematic of paradigms for MM stimulations with single 0.2 or 1U Col injection and repetitive daily vibration (2×3min) over MM. **(C)** The development of masticatory myalgia at an area over MM after single 1U Col injection into MM and daily vibration over MM area in male mice. Statistics is 2-way ANOVA (** p<0.01; n=5-7). **(D, E)** The development of headache-like behavior after a single 0.2U Col injection and daily vibration over MM area in male (*the panel D*) and female (the *panel E*) mice. Statistics is 2-way ANOVA (* p<0.05; ** p<0.01; n=5-7).

### Referred pain after eliciting masticatory myalgia from temporal muscle

TM stimulation and functional overload could contribute to myogenous TMD [12; 14; 43; 44]. Accordingly, we mimicked masticatory myalgia by stimulating TM with Col. After stimulation of TM with single injection of 10U Col (*Fig 4A*) or multiple low dose (0.2-0.5U) injections of Col (*Fig 4B*), mechanical hypersensitivity was assessed over MM at the same ipsilateral side relatively to TM injections and periorbital area (*Fig 1A*; green and red arrow area). Single 10U Col injection into TM generated myalgia in MM area for up to 12 days (*Fig 4C*) and substantial headache-like behavior up to 9 days in all tested male mice (*Fig 4D*). Repetitive low dose Col produced a more long-lasting masticatory myalgia in the MM area (*Fig 4C*), but not as persistent as during repetitive MM stimulations (*Fig 1E*). This TM stimulation paradigm also produced chronic headache-like behavior in male mice (*Fig 4D*). We next evaluated effects of TM stimulation by Col on masticatory myalgia and headache-like behavior in female mice. Single stimulation by higher dose Col produced significantly more myalgia in MM area for females compared to males (2-way ANOVA; F (10, 94) = 12.77; P<0.0001; n=6-7; *Figs 4C vs 4E*). Surprisingly, repetitive stimulations of TM triggered more pronounced myalgia in males versus females (2-way ANOVA; F (14, 149) = 6.317; P<0.0001; n=7; *Figs 4C vs 4E*). Single TM stimulation with higher dose of Col also generated drastically longer headache-like behavior in females compared to males (2-way ANOVA; F (10, 94) = 8.803; P<0.0001; n=6-7; *Figs 4D vs 4F*). In contrast, repetitive TM stimulations again revealed longer headache-like behavior in males (2-way ANOVA; F (15, 158) = 6.762; P<0.0001; n=7; *Figs 4D vs 4F*). Overall, this set of experiments demonstrated that modeling masticatory myalgia via TM stimulation by different paradigms induces pronounced referred pain and shows sex-dependency in myalgia and headache-like chronicity.

**Figure 4.**
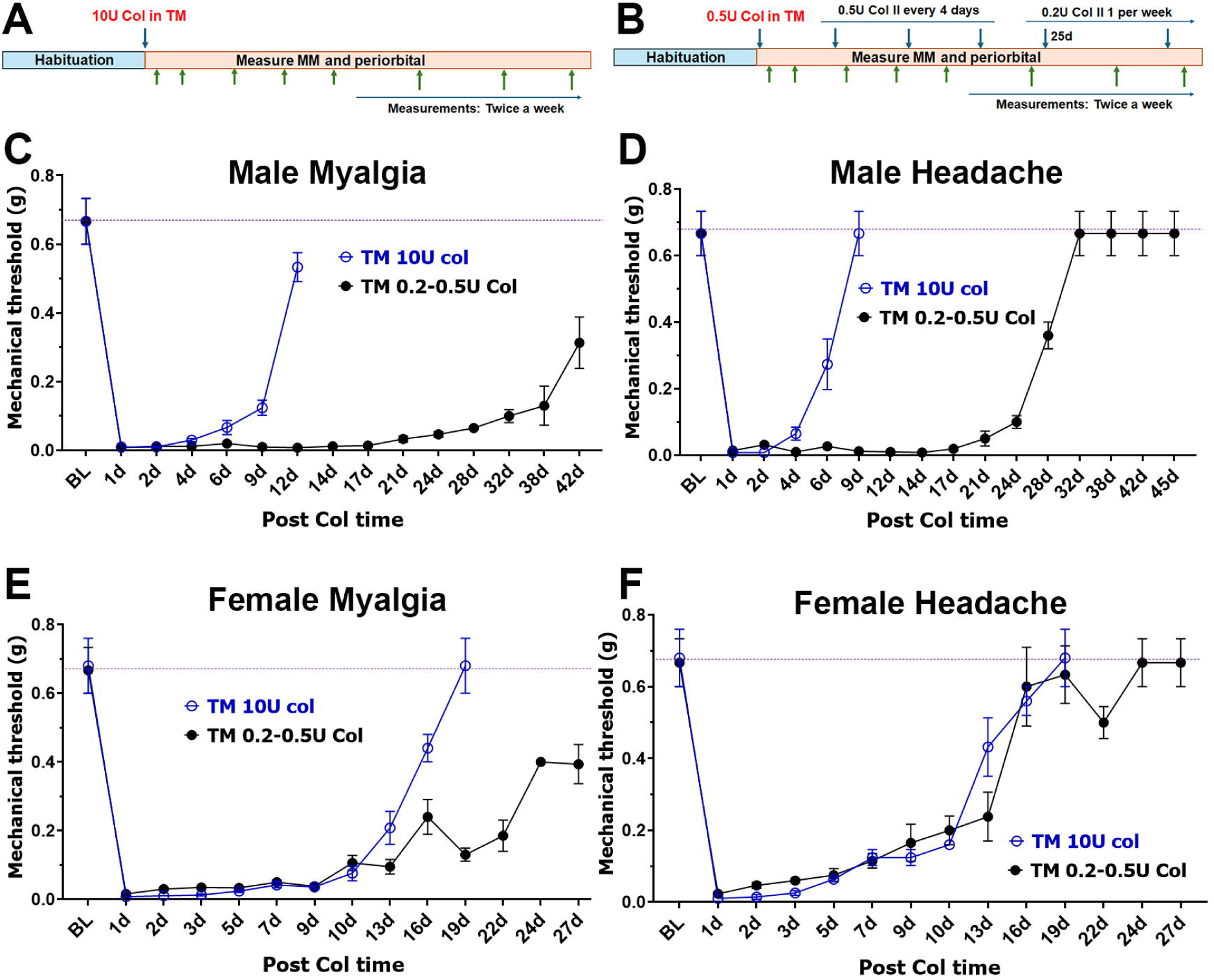
Referred pain after modeling masticatory myalgia by stimulating temporal muscle with collagenase type II. **(A, B)** Schematic of paradigms for TM stimulations with single 10U Col (*the panel A*) injection and repetitive 0.2-0.5U Col injections (*the panel B*). **(C, E)** The development of masticatory myalgia at an area over MM after single 10U Col injection into TM of male (*the panel C*) and female (*the panel E*) mice. **(D, F)** The development of periorbital mechanical hypersensitivity after a single 10U Col injection into TM of male (*the panel D*) and female (*the panel E*) mice.

### Referred pain after eliciting masticatory myalgia by forceful mouth stretch

It is believed that masticatory muscle overload could lead to myalgia [4; 48]. One of the approaches to model such overload is to perform forceful mouth stretches [7; 40]. We have measured mechanical hypersensitivity at MM area and periorbital area (*Fig 1A*, green and red arrows). It could be noted that beside modeling myogenous TMD, mouth stretch could also affect TMJ [45; 49]. In this respect, it is almost impossible to distinguish myogenous and arthragenous TMD in mice using the von Frey approach. Mouth stretch paradigms were implemented by several variants (*Fig 5A*). Mechanical hypersensitivity over MM was similar after single 45 min-long mouth stretch or 4 consecutive days by 45 min-long stretches (*Fig 5B*). A decrease from the 45 min-single stretch duration to 10 or 15 min reduced the length (in days) of mechanical hypersensitivity compared to single prolonged or repetitive stretches (*Figs 5B vs 5C*). Based on these results, we applied repetitive (every 5 days) 10 min-long stretch and added repetitive (3 days) restrain stress on the 11^th^ day as described [30] (*Fig 5A*). Such modeling produced persistent myalgia, which was similar for male mice with or without stress for first 20 days (*Fig 5D*). This mouth stretch paradigm produced short lasting (1 day) headache-like behavior in 66% (8 of 12) of the mice (*Fig 5E*). Mice undergoing repetitive mouth stretches did not exhibit headache-like behavior beyond 1d time point (*Fig 5E*). However, as expected, adding stress to mouth stretch paradigm elicited long lasting headache-like behavior starting from 14^th^ day post-first stretch (*Fig 5E*) [30]. Altogether, forceful mouth stretch modeling myogenous TMD, which likely induces arthragenous TMD as well, produced only 1 day-lasting and mild headache-like behavior in male mice.

**Figure 5.**
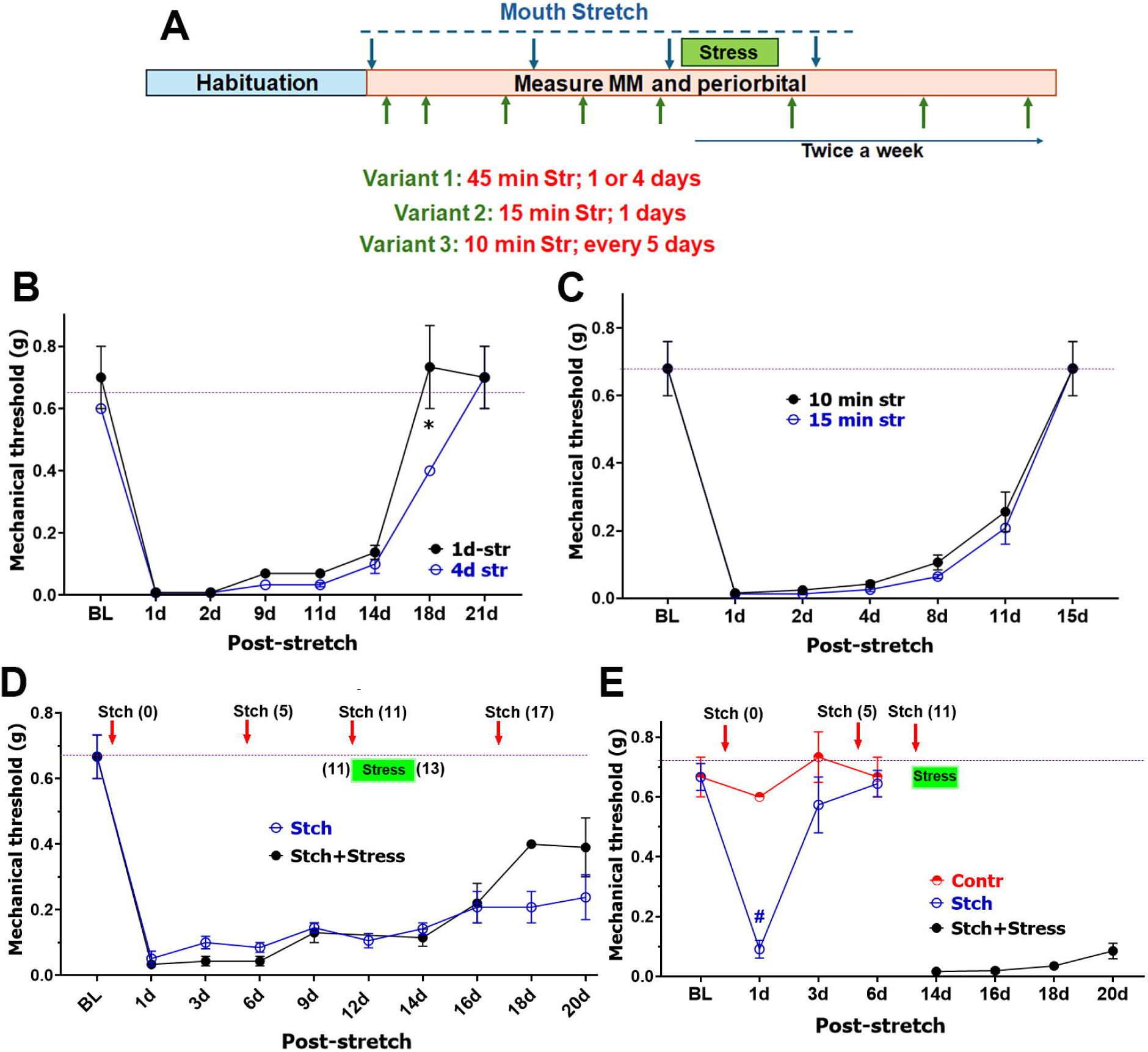
Referred pain after modeling masticatory myalgia by mouth stretch. **(A)** Schematic of induction of masticatory myalgia with mouth stretch and repetitive stress. **(B)** The development of masticatory myalgia at an area over MM after a single 45 min-long mouth stretch and 45 min-long mouth stretches 4 consecutive days in male mice. Statistics is 2-way ANOVA (* p<0.05; n=6). **(C)** The development of masticatory myalgia at area over MM after single 10 or 15 min-long mouth stretch in male mice. **(C)** The development of masticatory myalgia at area over MM after repetitive (every 5^th^ day) 10 min-long mouth stretch with additional (1h x 3) repetitive restraint stress between 11 and 14 days post first mouth stretch in male mice. **(D)** The development of headache-like behavior by repetitive (every 5^th^ day) 10 min-long mouth stretch with additional (1h x 3) repetitive restraint stress between 11 and 14 days post first mouth stretch in male mice. Statistics is 2-way ANOVA (# p<0.0001; n=6-12).

### Local referral gene plasticity after stimulation of temporal muscle

It is now known that referred pain have both central and peripheral mechanisms [42]. However, it is unknown whether pain referral is accompanied by local gene plasticity at the referred pain sites. Thus, we examined whether stimulation of the TM led to gene plasticity in MM and dura mater. According to observed headache-like referred pain elicited upon TM stimulation by single injection of 10U Col in males and females (*Figs 4D, 4F*), we isolated dura mater (n=3-6) at 2d post Col injection (n=6). Dura mater isolated from male and female mice injected with saline into TM served as controls (n=3-4). Transcriptomic changes in dura mater were evaluated using bulk RNA-seq [26]. Selection criteria for differentially expressing gene (DEGs) were FC>1.5; FPKM>1 and Padj<0.05. In males, 439 DEGs were up and 445 DEGs were down-regulated in dura mater with FC>2, and 1013 DEGs were up and 1019 DEGs were down-regulated in dura mater with FC>1.5. In females, 880 DEGs were up and 549 DEGs were down-regulated in dura mater with FC>2, and 1522 DEGs were up and 1321 DEGs were down-regulated in dura mater with FC>1.5. Tissue repair and morphogenesis, axonogenesis, gliogenesis, synaptogenesis, and neurogenesis processes dominated up-regulated DEGs at 2d post Col in male and female dura mater (*Figs 6A, 6B*). Difference in transcriptomic changes between males and females were minor. Thus, male had gliogeneses, while females showed processes related to synaptogenesis and neurotransmission (*Figs 6A, 6B*). The major tissue undergoing morphogenesis was muscle in females, while processes related to bone remodeling were additionally present in males (*Figs 6A, 6B*). Immune system related processes dominated down-regulated DEGs in male and female dura mater (*Figs 6C, 6D*). Again, the observed differences were minor and related to presence of skin development processes in males, while females exhibited more pronounced down-regulation of immune system related DEGs in males (*Figs 6C, 6D*). Comparisons for biological processes between males and females using PANTHERdb software did not yield any processes for both up- and down-regulated DEGs. Despite similarities in biological processes undergoing plasticity in dura mater of males versus females, actual overlaps of DEGs were low, 15.5% for up- and 13.9% for down-regulated DEGs (*Figs 6E, 6F*).

**Figure 6:**
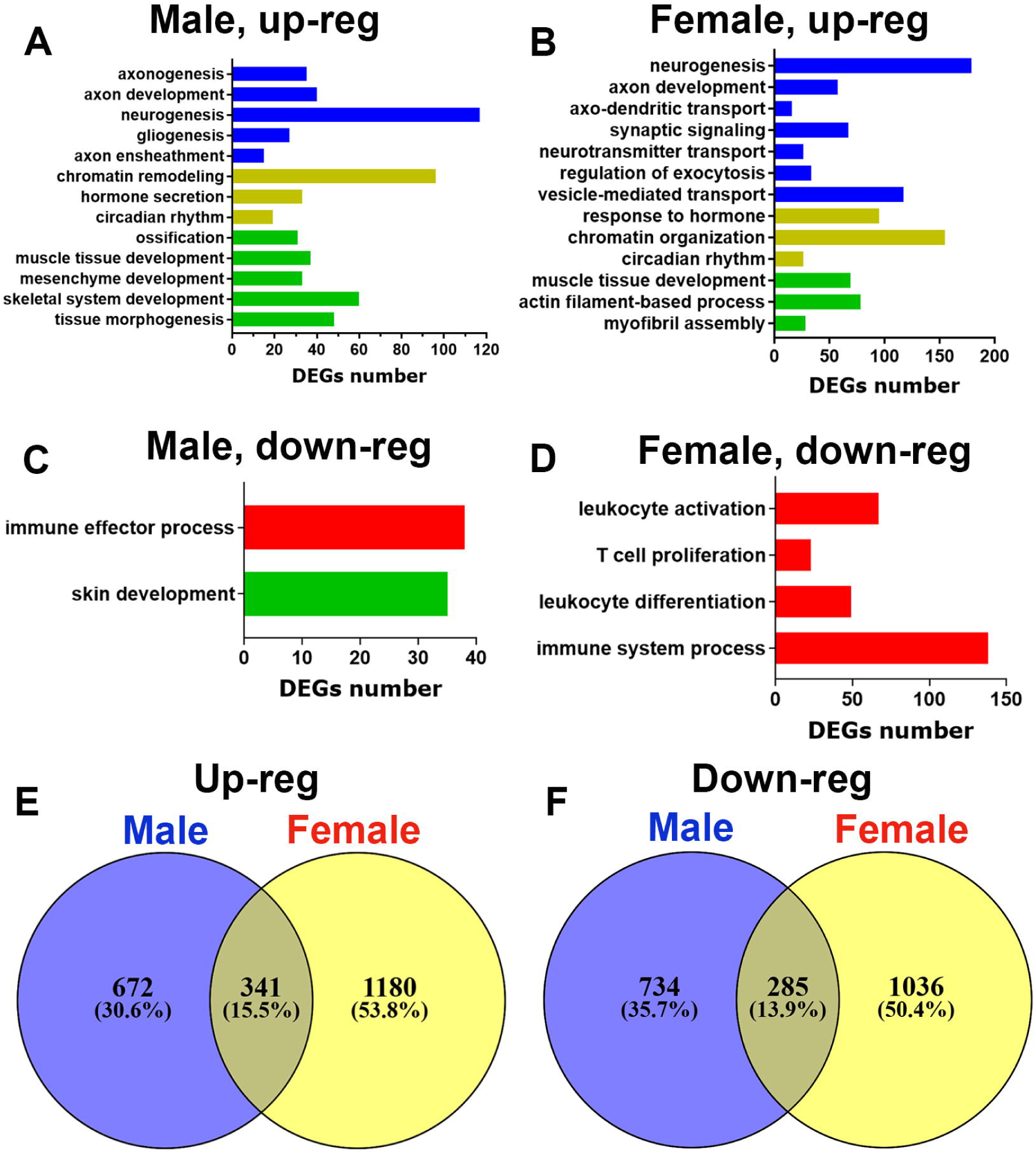
Up- and down-regulated biological processes in the dura mater after single Col injection into TM of male and female mice. (**A, B**) Biological processes for up-regulated DEGs in the dura mater at 2d post-Col (10U) into TM of male (*panel A*) and female mice (*panel B*). (**C, D**) Biological processes for down-regulated DEGs in the dura mater at 2d post-Col (10U) into TM of male (*panel C*) and female mice (*panel D*). (**E**) Venn diagrams of 10U Col-induced up-regulation of DEGs in the dura mater at 2d post-TM treatments for males versus females. (**F**) Venn diagrams of 10U Col-induced down-regulation of DEGs in the dura mater at 5d post-TM treatments for males versus female mice. The X-axis on *the panels A-D* represents the number of DEGs. The Y-axis notes biological processes.

Stimulation of TM by repeated application of low dosages of Col generate chronic mechanical hypersensitivity with resolution in MM of female mice (*Fig 4E*). Resolution phase of MM Col injection-induced masticatory myalgia is accompanied by sharp up-regulation of immune system related DEGs [26]. Accordingly, we investigated whether repetitive stimulation of TM with low doses of Col (*Fig 4B*) leads to gene plasticity in MM from females during resolution phase (28 days post-first Col injection; *Fig 4E*). This treatment led to up-regulation of 234 DEGs and down-regulation of 177 DEGs in MM with FC>2; and 330 DEGs were up and 259 DEGs were down-regulated in MM with FC>1.5. Statistically significant up-regulated biological process revealed by PANTHERdb software included developmental process in muscles and bones, extracellular matrix remodeling, gliogenesis, neurogenesis, synaptogenesis and extensive changes in expressions of genes related to the immune system (*Fig 7A*). In contrast, few biological processes underwent down-regulation in MM (*Fig 7B*). Interestingly, up- and down-regulated processes in MM during resolution phase were similar when Col was injected into TM versus MM [26]. Thus, key processes such as extracellular matrix remodeling, synaptogenesis, neurogenesis, gliogeneses and especially, escalation of inflammatory processes were present whether Col was administrated into MM or TM. In summary, referred pain in head and neck area is associated with local gene plasticity at the site to which mechanical hypersensitivity was referred.

**Figure 7:**
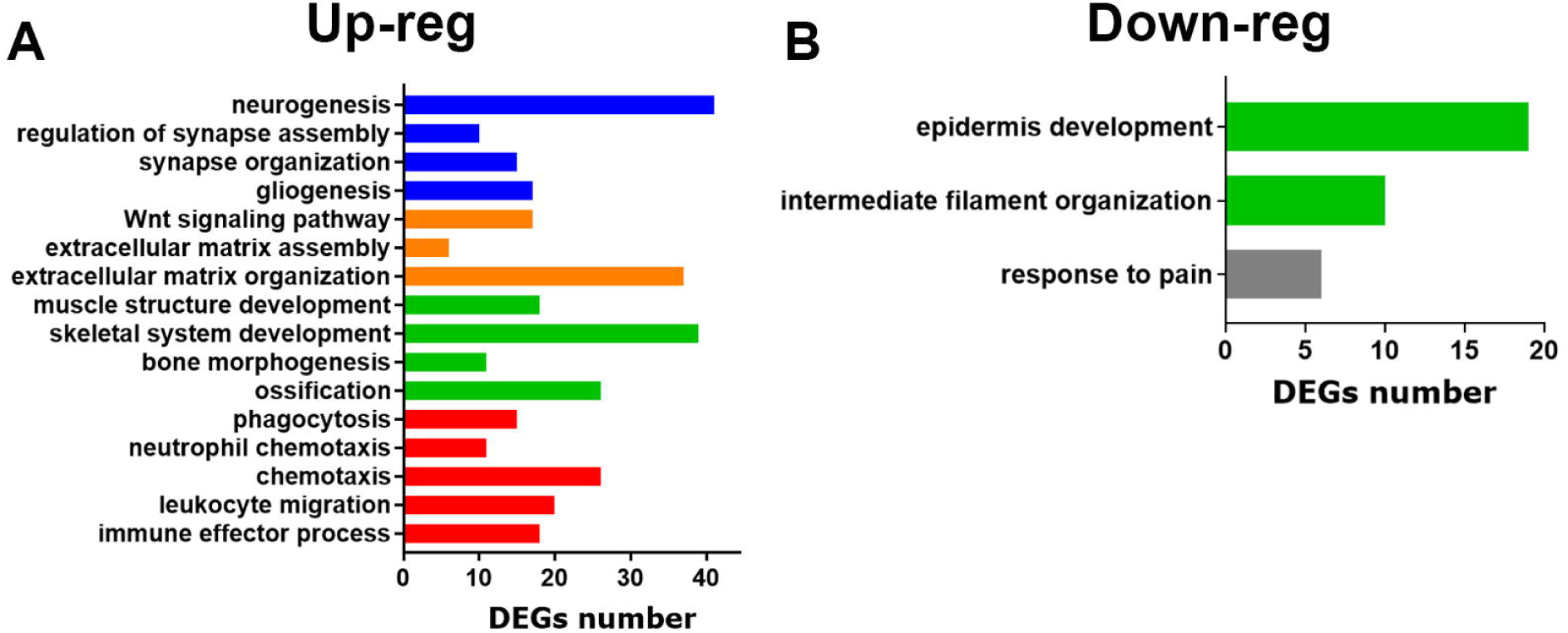
Up- and down-regulated biological processes in the MM after repetitive Col injections into TM of female mice. (**A, B**) Biological processes for up-regulated (*the panel A*) and down-regulated DEGs (*the panel B*) in the MM at 28d post-Col treatment of TM of female mice. The X-axis on *the panels A-B* represents the number of DEGs. The Y-axis notes biological processes.

## Discussion

A substantial subset of patients with myogenous TMD (TMD-M) report widespread and referred pain, including headache, pain within TMJ and neck [5; 29; 31]. Clinical data indicates that both peripheral and central mechanisms play roles in referred pain in the head and neck area [10; 13; 16; 19; 42]. However, intricate and detailed mechanisms and neuronal circuitries involved in referred pain are largely unknown. Hence, the need for more clinically accurate TMD-M animal models is crucial in order to study these intricate mechanisms explaining referred pain. Accordingly, this study aimed to model referred pain in the head and neck area in animals; to investigate whether main masticatory muscles - temporal and masseter - have distinct contributions to referred pain; and to understand whether local mechanisms are restricted only to pain eliciting site.

TMDM is thought to develop due to muscle ischemia following consistent and repetitive physical overloading of masticatory muscles [15; 28]. Ischemia-induced muscle damage involve extracellular matrix (ECM) dysfunction/damage [6; 23; 46] and collagen degradation [21; 24]. To mimic these clinical aspects of TMD-M, we modeled masticatory myalgia by different means, Including single and repetitive Col injections, administrations of low and high dosages of Col, vibration-based stimulation of an area over masticatory muscles, and forceful stretching of mouth.

It appear that stimulation of TM, but not MM, produced reliable, substantial, longer-lasting and sex-dependent referred pain in a head and neck area, including headache. Nevertheless, repetitive mild vibration over MM produced a slight referred pain - headache. In clinical setting, expansions of receptive fields within muscle area are often denoted as “referred pain”, which is routinely observed for MM area [10; 19]. Referred pain from facial muscles to headache, or from TM to MM requires more complicated neuronal circuitry. Thus, MM is innervated by masseteric nerve (MN) and TM by posterior deep temporal nerve (TN). Both MN and TN are branched from inferior alveolar nerve originated in the trigeminal ganglion (TG) V3 region [11], while dura mater is innervated by TG neurons of ophthalmic branch (V1). Additionally, some studies indicated that MN and TN could also innervate tissues surrounding the TMJ condyle [11]. Referred pain is likely due to sensitization of second-order brain stem neurons, which are convergence points for sensory neuron afferents innervating different parts of the head and neck area [10; 19]. In this respect, the anatomical arrangements of nerves imply that referred pain from TM to MM and vice versus, as well as from MM or TM to TMJ could be relatively easier to achieve than referred pain from masticatory muscles to headache. Moreover, limited information on the neuronal circuitry architecture for TG nerve projections into brain stem does not allow clear explanation on why TM, but not MM, stimulation trigger headache. One possibility is that stimulation of TM “spills” to skin over TM, which is innervated by TG V1 neurons. However, our pilot experiments with methylene blue injection into TM showed that only 5-10% of diffusion from TM to skin occurred. Another possibility is that TM and dura mater innervating TG neurons spatially closely located within the TG, despite they belong to V1 and V3 areas. Such close locations are possible, since boundaries between V1, V2 and V3 TG regions are widely dispersed. Overall, much work need to be done to clearly understand converging neuronal circuitries connecting TN (and MN) with dura mater nerve projections in the brain stem nuclei.

Contributions of local (aka peripheral) mechanisms to referred pain is well documented [13; 42]. These mechanisms suggest sensitization of neurons/nerves innervating sites from which referred pain was originated. Our data implied that local sites to which pain was referred could be substantial contributors to the referral pain phenomenon in the head and neck area. Thus, stimulation of TM led to gene plasticity in MM. Moreover, TM-induced gene plasticity in MM was similar to those observed after direct stipulation of MM [26]. Gene plasticity in dura mater was not characterized for different headache models. However, our unpublished data suggest that TM stimulation-induced gene plasticity in dura mater resembles to expression changes detected in dura mater during headache triggered by irregular sleep pattern. In summary, these finding could suggest an alternative mechanism for referral pain. According to this mechanism, referral pain phenomenon could be due to sensitization of nerves innervating multiple local sites, which creates a peripheral network accounting for spreading pain in the head and neck area.

## Conflict of interest statement

The authors have no conflicts of interest to declare.

## Acknowledgments

We would like to thank Dr. Gregory Dussor (UT Dallas; Richardson, Texas) for his guidance in setting up measurements of mechanical hypersensitivity in orofacial region, including periorbital area. We would also like to thank Mrs. Korri Weldon from Dr. Zhao Lai’s lab (UTHSCSA; San Antonio, Texas) for performing bulk RNA sequencing procedure on all samples.

This research work was supported by HEAL Initiative NIDCR/NIH DE029187 (to A.N.A.); by the National Institute Of Arthritis And Musculoskeletal And Skin Diseases of the National Institutes of Health (NIH/NIAMS) through the NIH HEAL Initiative (https://heal.nih.gov/) The Restoring Joint Health and Function to Reduce Pain (RE-JOIN) Consortium UC2 AR082195 (to A.N.A.); by Summer Physiology Undergraduate Researcher (SPUR) Program R25 NS 115552 (to J.A.) and by Co-STAR training grant 2T32 DE014318-22 (to J.A.).

## Author Contributions

Conceptualization: M.E., M.D.B, K.R. and A.N.A.; Methodology: A.H.H, J.A., J.M.M., F.A., W.G., and J.Y.; Formal Analysis: A.H.H, J.A., J.A., Y.Z., Z.L. and A.N.A.; Investigation, J.A., J.A, W.G., and J.Y.; Writing—Original Draft: A.N.A.; Writing—Review & Editing, J.A., Z.L., M.E., M.D.B, K.R. and A.N.A.

## Data availability statement

RNA-seq data has been deposited to GEO Accession, the number is pending (GSExxxx).

## Notes

### Competing Interest Statement

The authors have declared no competing interest.

